# Human norovirus efficiently replicates in differentiated 3D-human intestinal enteroids

**DOI:** 10.1101/2022.06.09.495585

**Authors:** Carmen Mirabelli, Nanci Santos-Ferreira, Merritt G. Gillilland, Roberto J. Cieza, Justin A. Colacino, Jonathan Z. Sexton, Stefan Taube, Joana Rocha-Pereira, Christiane E. Wobus

**Affiliations:** Institute for Virology and Cell Biology, University of Lübeck, Germany; Department of Microbiology and Immunology, University of Michigan, Ann Arbor, MI, United States of America; KU Leuven, Department of Microbiology, Immunology & Transplantation, Rega Institute, Laboratory of Virology & Chemotherapy, Leuven, Belgium; Department of Internal Medicine, University of Michigan, Ann Arbor, MI, United States of America; Department of Environmental Health Sciences, University of Michigan, School of Public Health, Ann Arbor, MI, United States of America; College of Pharmacology, University of Michigan, Ann Arbor, MI, United States of America

**Keywords:** Human norovirus, 3D-human intestinal enteroids, polarity inversion, RNA sequencing

## Abstract

Human norovirus (HNoV) accounts for one fifth of all acute viral gastroenteritis worldwide and an economic burden of ∼$60 billion globally. The lack of treatment options against HNoV is in part due to the lack of cultivation systems. Recently, a model of infection in biopsies-derived human intestinal enteroids (HIE) has been described: 3D-HIE are first dispersed in 2D-monolayers and differentiated prior to infection, resulting in a labor-intensive, time-consuming procedure. Here, we present an alternative protocol for HNoV infection of 3D-HIE. We found that 3D-HIE differentiate as efficiently as 2D-monolayers. In addition, immunofluorescence-based quantification of UEA-1, a lectin that stains the villus brush border, revealed that over 90% of differentiated 3D-HIE spontaneously undergo polarity inversion, allowing for viral infection without the need for microinjection. Infection with HNoV GII.4-positive stool samples attained a fold-increase over inoculum of ∼2 Log_10_ at 2 days post infection or up to 3.5 Log_10_ when ruxolitinib, a JAK1/2-inhibitor, was added. Treatment of GII.4-infected 3D-HIE with the polymerase inhibitor 2’-*C*-Methylcytidine (2CMC), other antivirals, or with a HNoV-neutralizing antibody showed a reduction in viral infection, suggesting that 3D-HIE are an excellent platform to test anti-infectives. The host response to HNoV was then investigated by RNA sequencing in infected versus uninfected 3D-HIE, in the presence of ruxolitinib to focus on viral-associated signatures. The analysis revealed upregulated hormones and neurotransmitter signal transduction pathways and downregulated inflammatory pathways upon HNoV infection. Overall, 3D-HIE have proven to be a more robust model to study HNoV infection, screen antivirals and investigate host response to HNoV infection.

**Importance:** Human norovirus (HNoV) clinical and socio-economic impact calls for immediate actions in the development of anti-infectives. Physiologically-relevant *in vitro* models are hence needed to study HNoV biology, tropism and mechanism of viral-associated disease but also as a platform to identify antiviral agents. Biopsy-derived human intestinal enteroids are a biomimetic of the intestine and recently described as a model that supports HNoV infection. The established protocol is time-consuming and labor-intensive. Therefore, we sought to develop a simplified and robust alternative model of infection in 3D enteroids that undergo differentiation and spontaneous polarity inversion. Advantages of this model are the shorter experimental time, better infection yield and spatial integrity of the intestinal epithelium. This model is potentially suitable for the study of pathogens that infect intestinal cells from the apical surface but also for unraveling the interactions between intestinal epithelium and indigenous bacteria of the human microbiome.

## Introduction

Diarrheal diseases are the fourth cause of death worldwide and the second cause of morbidity in children less than 5 years old(1). In particular, human norovirus (HNoV) is the main causative agent of viral gastroenteritis worldwide, with a clinical burden of nearly 200,000 hospitalizations in Europe(2) and an economic burden of 65 billion US dollars, worldwide(1). The virus has been poorly characterized in terms of its viral life cycle, tropism, and pathophysiology due to the lack of easy-to-use cell culture systems that support robust viral replication and ensure the production of cell culture-derived viral stocks. Consequently, the development of prophylactic and therapeutic agents has been challenging. The human intestinal enteroid culture (HIE), a biopsy-derived model of the intestinal epithelium, is an emerging tool to study enteric viruses that have been refractory to transformed cell culture systems. It proved efficacious to support the cultivation of a wide range of enteric pathogen, including HNoV(3). Drawbacks of this systems are the time-consuming nature of the infection protocol, the elevated costs for the maintenance of the culture, the lack of long-term passage of HNoV, and the inability to generate a high-titer viral stock. The published protocol of infection includes the dispersion of HIE from 3D spheres into 2D monolayer on collagen-coated plates(3). For one well of a 96-well plate, it is recommended to use 100,000 cells, 10X more than transformed cell lines. Two to six days later, when HIE 2D-monolayers reach confluency, cells are differentiated by withdrawal of Wnt3a for 5-6 days before infection is initiated. Thus, the experimental time from HIE splitting to infection is 15 days, on average. Another disadvantage to the use of HIE 2D monolayers is the loss of 3D spatial organization, that is crucial for intestinal physiology(4). For all these reasons, we sought to investigate whether differentiated 3D-HIE are amenable to HNoV infection. However, 3D-HIE are organized with the apical surface of the cells towards the lumen of the sphere, therefore the exposure to a pathogen might be limited. Recently, a protocol to invert HIE polarity was published (**Apical-Out** HIE)(5),whereby the removal of basal membrane extract (BME) ensures polarity reversal. The mechanism is dependent on extracellular matrix protein (ECM) concentration, because addition of ECM to HIE in the absence of BME blocks polarity reversion(5). Apical-Out HIE thus provide a potential solution to ensure greater accessibility of 3D-HIE to pathogens.

The induction of the host innate response plays an essential role in the suppression of pathogen infection. In the case of enteric pathogens, interferon responses are highly upregulated *in vivo* and *in vitro*(6, 7). We previously reported that human astrovirus VA1, another enteric virus and causative agent of viral gastroenteritis, induces interferon and interferon-stimulated genes (ISG) in the HIE system but not in an immortalized cell line, the Caco2 cells(8). This also suggest that, intrinsically, the HIE culture restricts enteric virus infection more efficiently than immortalized cell lines. For this reason, a JAK inhibitor, ruxolitinib, has been successfully added to the infection protocol to prevent stimulating the innate immune response and thus increase the yield of HNoV in HIE(9).

Here, we describe the establishment of a HNoV infection protocol in 3D-HIE. We characterized the differentiation state of 3D-HIE compared to 2D-HIE and the degree of polarity inversion in 3D-HIE upon differentiation. We demonstrate that 3D-HIE are amenable to infection with HNoV with a reduction of experimental time (9 vs 15 days) and increased yield (fold increase over inculum) and reproducibility. We also characterized the host response of 3D-HIE to HNoV infection in culture treated with ruxolitinib by RNA sequencing. Altogether, we describe an adapted protocol for HNoV infection in 3D-HIE that is amenable to antiviral discovery and virus-host interaction studies. In addition, this model may facilitate future studies of other enteric pathogens/microbes that infect the apical surface of the intestinal epithelium.

## Results

### 3D-HIE undergo terminal differentiation similar to 2D-HIE, but infection with HNoV results in better yields

Differentiation of HIE is achieved by withdrawal of Wnt3a from the maintenance media, that triggers the development of a heterogeneous, terminally differentiated epithelium by day 6(8). We sought to determine if HIE derived from fetal ileum HT124 undergo terminal differentiation when maintained in 3D in basal membrane extract (BME). Towards that end, we monitored the mRNA levels of differentiation markers by RT-qPCR. As a control, 2D-monolayers of HT124 HIE were also prepared and differentiated in collagen-coated plates. At 6 days post differentiation, comparison of mRNA levels of Lgr5, lysozyme, mucin 2 (MUC2) and sucrase isomaltase (SI) as markers of stem-cells, Paneth cells, goblet cells and mature enterocytes, respectively, revealed similar transcript levels in differentiated 2D *versus* 3D-HIE (Figure 1A). This suggests that the withdrawal of Wnt3a triggers terminal differentiation of 3D-HIE in BME to the same extent and in the same time frame as 2D-HIE. Representative images of differentiated 2D- and 3D-HIE are shown in Figure 1B. To address whether differentiated 3D-HIE could support HNoV infection, GII.4-positive stool samples were used to infect the 3D-HIE, after removal from BME. After a 2 h incubation at 37°C in the presence of the bile acid GCDCA (500µM), the inoculum was washed off, one set of 3D-HIE was harvested in TRI Reagent (2 h), while another set was re-embedded in BME and kept at 37°C for 2 days (2 d) in differentiation media supplemented with GCDCA (500µM) and a JAK inhibitor ruxolitinib (2µM). Differentiated 2D-HIE were infected in parallel according to the published protocol, also in the presence of GCDCA (500µM) and ruxolitinib (2µM). Quantification of HNoV yields by RT-qPCR revealed a 2.5 Log_10_ versus 1 Log_10_ of fold increase in 3D-HIE versus 2D-HIE, that was achieved over only 2 d post infection (dpi) (Figure 1C). These data suggest that 3D-HIE differentiation is sufficient to support HNoV infection, and that in our system, the protocol of infection of 3D-HIE results in better yield (fold increase over inoculum) as compared with the established 2D-HIE protocol.

**Figure 1:**
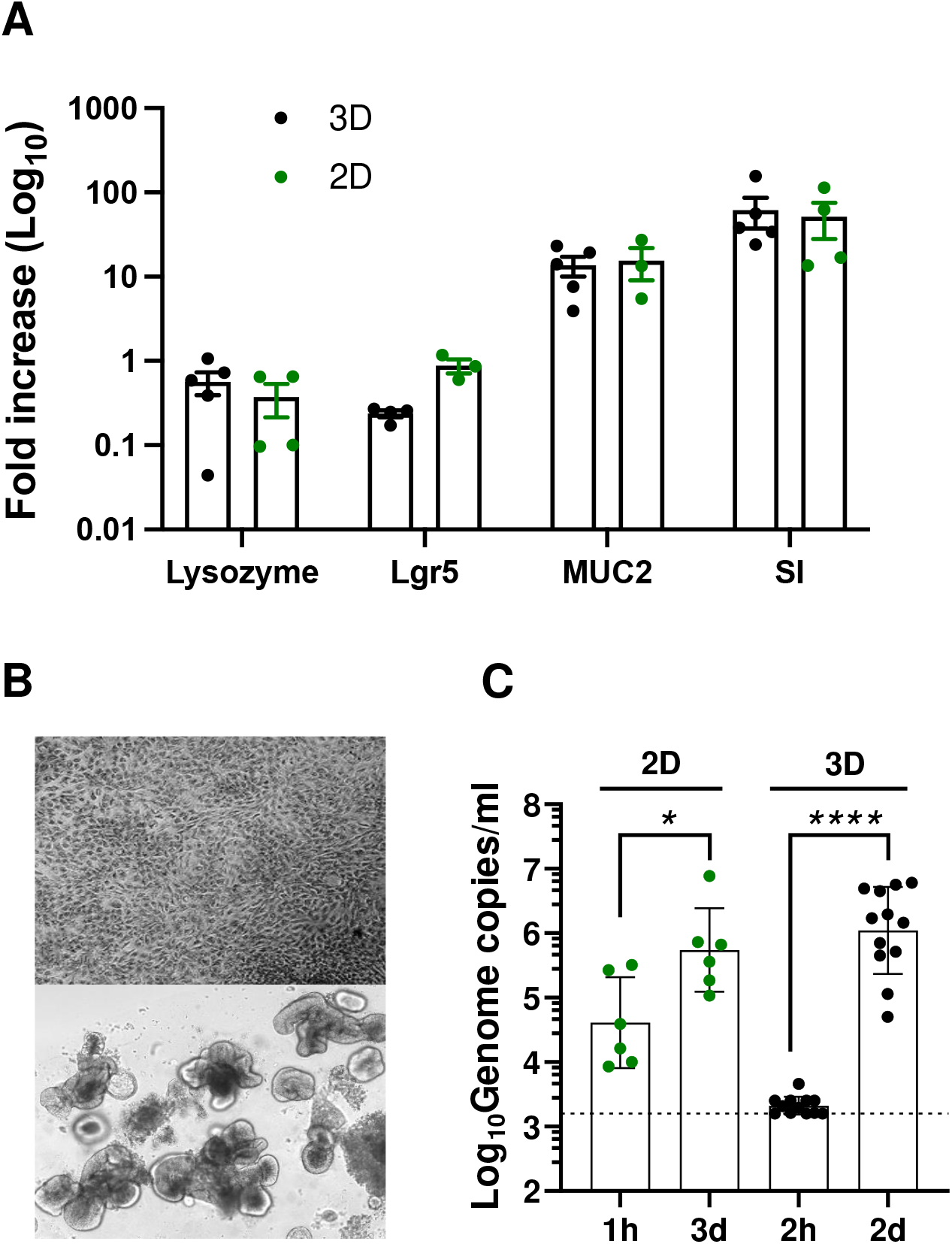
3D-HIE undergo terminal differentiation and support HNoV replication. A) Differentiation of 3D-HIE (3D), compared with 2D-HIE monolayers (2D). HIE from a fetal donor HT124 have been differentiated by Wnt3a removal in 3D spheres or after dispersion in a collagen-coated plate for 6 days. mRNA levels of selected host genes were quantified by qPCR. Lysoyzme: marker of Paneth cells, Lgr5: marker for crypt (stem)-cells, mucin 2, MUC2: marker of goblet cells and sucrase isomaltase, SI: marker for mature enterocytes. Each dot represents a biological replicate. B) Representative images of differentiated HT124 2D- and 3D-HIE. C) 2D and 3D-HIE were infected with HuNoV GII.4-positive stool sample (#233). 2D-HIE were infected according to the published protocol in the presence of ruxolitinib (2µM) and harvested at 3dpi. 3D-HIE were removed from BME and infected for 2hrs at 37°C. After, 3D HIE were washed twice and one set was harvested in TRI Reagent (2h) and another set was seeded in BME and maintained in differentiation media with GCDCA (500µ) and ruxolitinib (2µM). Two days post infection, 3D HIE were harvested in TRI Reagent (2d), RNA was extracted from all the samples and RT-qPCR was used to establish viral titers. Each dot represents an independent biological replicate (independent infection).*p value <0.05,****p value <0.0001, student t-test calculated in GraphPad Prism.

### Differentiation of 3D-HIE induces a spontaneous inversion of polarity that grants HNoV accessibility

Efficient infection of 3D-HIE by HNoV suggest that the virus has access to the apical surface of enterocytes. Interestingly, 3D-HIE have been described to revert polarity spontaneously after removal of BME. The inversion is dependent on the percentage of extracellular matrix (ECM) proteins in the media(5). Because our differentiation protocol starts 3-4 days after passaging, resulting in 3D-HIE being embedded in BME for ∼9-10 days, we hypothesized that the consequent reduction in % ECM proteins (caused by medium changes) could trigger spontaneous inversion. We also sought to determine whether the process of terminal differentiation, with Wnt3a withdrawal, may further enhance the polarity inversion. To this end, we differentiated 3D-HIE for 6 days and as a control, we kept in parallel 3D-HIE in proliferation media (Figure 2A). Next, differentiated (-Wnt3A) and undifferentiated (+Wnt3a) 3D-HIE were removed from BME, fixed with paraformaldehyde (4% in PBS-/-) and stained with DAPI for nuclear detection and *Ulex Europeus Agglutinin-*1 (UEA-1), a lectin that binds the α-L-fucosyl residues of glycoproteins and glycolipids at the apical surface of the intestinal epithelium(8). For ease of analysis, 10-20 3D-HIE were transferred into a well of a poly-lysin-coated black 96-well plate and subjected to confocal microscopy imaging with the CellVoyager CQ1 high-content microscope (Yokogawa). Image segmentation by Cell Profiler and analysis of the UEA-1 positive 3D-HIE revealed that after 10 days of culturing in proliferation media, about 50% of 3D-HIE spontaneously inverted polarity. In differentiation media, the proportion of UEA-1-positive spheroids was ∼ 90% (Figure 2B). Because a protocol of polarity inversion was recently published(5), we next wanted to compare the efficacy of replication in 3D-HIE generated with the published protocol (3D-HIE without BME, **Apical-Out**) or with our protocol (3D-HIE embedded in **BME**). Interestingly, the efficacy of replication at 2 dpi was >2 Log_10_ higher in ECM 3D-HIE than in HIE generated with the Apical-Out protocol (Figure 3C). These data suggest that the apical surface of differentiated 3D-HIE is exposed and explain why these cultures are amenable to infection with HNoV without the burden of microinjection. In addition, BME appears to support infection by HNoV as **Apical-Out** HIE did not reach the same yield of infection.

**Figure 2:**
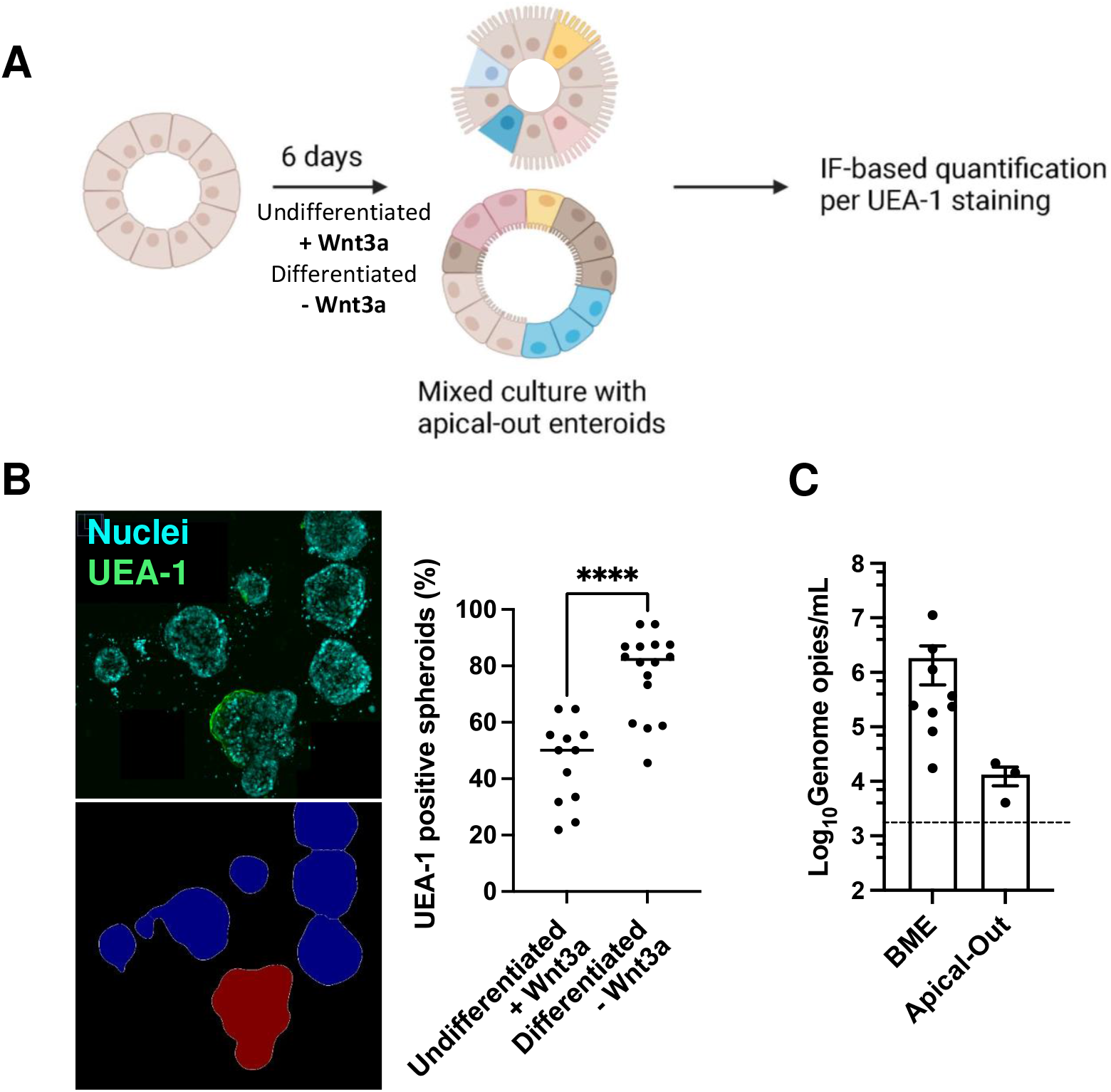
3D-HIE spontaneously revert polarity upon differentiation. A) Schematic of 3D-HIE differentiation and pipeline for quantification of 3D-HIE with inverted polarity, by immune fluorescence after staining with the lectin *Ulex Europaeus Agglutinin-1*, UEA-1. B) Representative image and image segmentation of 3D-HIE (left). 3D-HIE were cultured for 6 days with maintenance or differentiation media. The 3D spheres were then collected and seeded in a 96-well black collagen-coated plate. 3D-HIE were fixed for 20 minutes with 4% PFA and stained with UEA-1 and DAPI. Plates were imaged with the CellVoyager CQ1 scanning disk, automated, high content imaging microscope (Yokogawa). From each well, images from 16 fields were collected, covering ∼80% of the surface area with a 10X objective. The graph (right) represents quantification of of 3D-HIE spheres, positive for UEA-1 after image analysis with CellProfiler software. Each dot represents a technical replicate from 3 independent experiments. ****p value <0.0001, student t-test calculated in GraphPad Prism. C) 3D-HIE were first subjected to the apical-out protocol (Apical-Out), then differentiated and infected with GII.4-positive stool sample, #20942. In parallel, 3D-HIE were differentiated in BME and infected with #20942. Cells were harvested at 2dpi and viral genome copies were quantified by RT-qPCR. Each dot represents an independent biological replicate.

**Figure 3:**
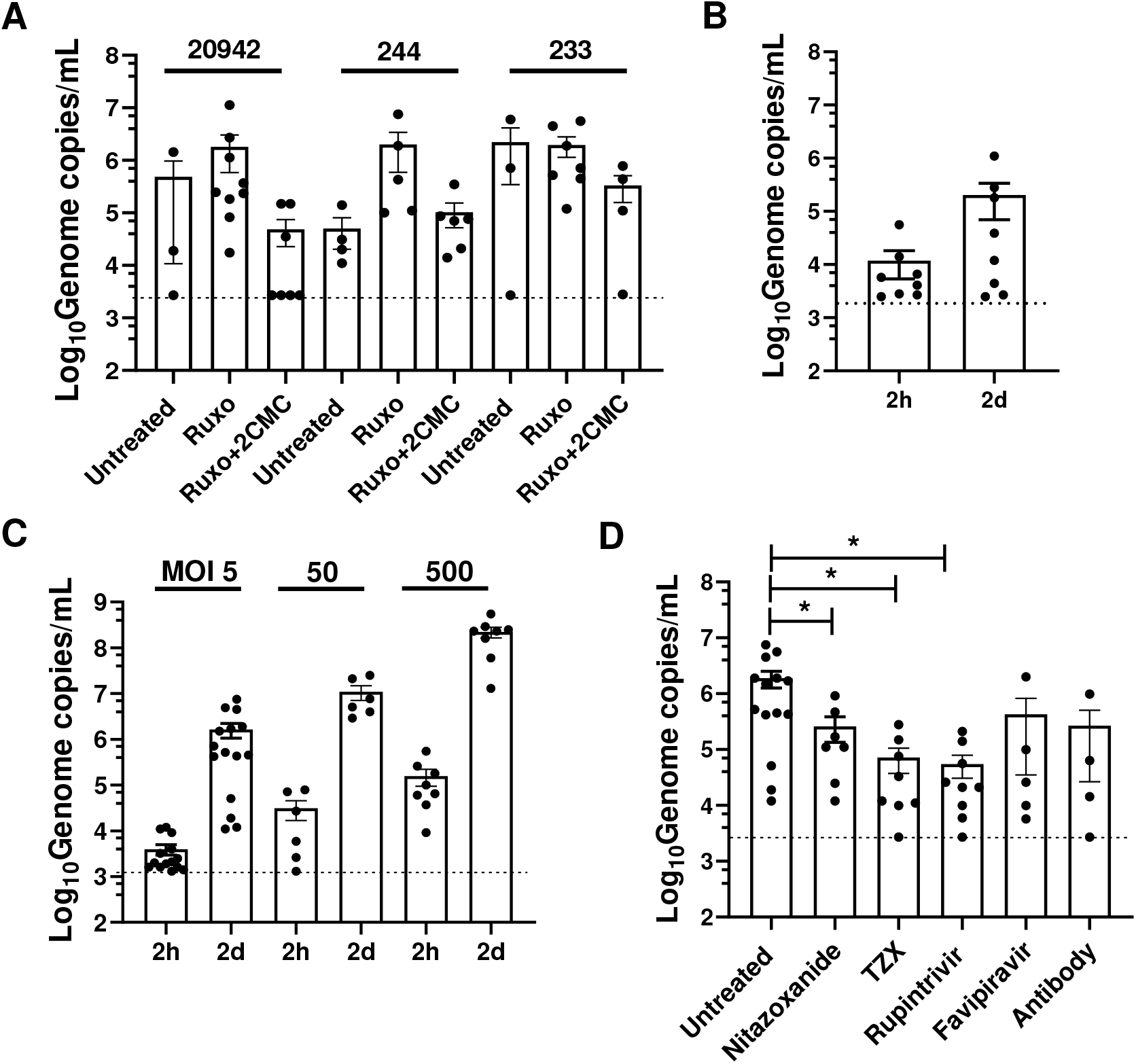
3D-HIE constitute a robust platform for HNoV infection and screening of anti-infectives. A) 3D-HIE were infected with filter stool positive for GII.4 (#233, 244 and 20942) and after washing, 3D-HIE were seeded in BME and maintained in differentiation media with GCDCA (Untreated), with ruxolitinib (2µM, Ruxo) or with the nucleoside inhibitor 2’-C-methylcytidine (50µM, Ruxo + 2CMC). Cells were harvested 2 dpi and viral genome copies were calculated by RT-qPCR. B) 3D-HIE derived from the J2 adult jejunum line were infected with stool positive for HNoV GII.4 (data from different strains was pulled together) and harvested at 2h and 2dpi. C) 3D-HIE were infected with increasing concentration (MOI) of HNoV GII.4 (data from different strains was pulled together) and were harversted at 2h and 2d for viral genome copies quantification by RT-qPCR. D) 3D-HIE were infected with HNoV GII.4 (data from different strains was pulled together) and treated with selected compounds, Nitazoxanide (50µM) and its active metabolite TZX (50µM), the enterovirus 3C-protease inhibitor rupintrivir (50µM), the polymerase inhibitor Favipiravir (250µM) and the neutralizing antibody (md145-22-1-2, R-biopharm). The small molecules antivirals were added after infection in differentiation media whereas the antibody was used to pre-treat the virus prior to infection for 30 minutes. For all graphs, each dot represents a technical replicate of at least 3 independent biologic experiments. Statistical analysis was performed with GraphPad Prism. *p-value <0.05, student t-test.

### HNoV infection of 3D-HIE is robust, dose-dependent and sensitive to antiviral treatment

To characterize the infection of 3D-HIE in further detail, we evaluated the efficacy of infection of a number of HNoV-positive stool samples in the presence or absence of ruxolitinib, a JAK inhibitor that limits interferon response and the nucleoside inhibitor 2’-*C*-Methylcytidine (2CMC). Replication of stool samples 20942 and 244, but not 233 was improved by adding ruxolitinib (Figure 3A), further substantiating the critical importance of the interferon response in restricting HNoV infection of HIE(9). In addition, treatment with the nucleoside inhibitor 2’-*C*-Methylcytidine (2CMC) reduced replication for all three stool samples at 2 dpi (Figure 3A). To confirm translatability of findings to other HIE, we also assessed HNoV replication in a HIE line derived from adult jejunal tissues (J2). We observed a 1 Log_10_ increase at 2 dpi versus 2 hpi (Figure 3B), suggesting that infection occurs in this adult line also in the 3D-format but the permissiveness of infection is reduced as compared to the fetal line HT124. We next infected 3D-HIE with increasing doses of HNoV-positive stool samples (Figure 3C). A dose-dependent increase of replication was observed at 2 dpi. However, the fold increase between 2 dpi and 2 hpi remained constant at ∼2.5 Log_10_, suggesting that the system supports a defined amount of infection regardless of the initial infectious dose.

To test whether the infection model could be used to test antivirals, we lastly infected 3D-HIE in the presence of compounds that were previously described as active against norovirus: Nitazoxanide, and its active metabolite TZX(10), the protease inhibitor Rupintrivir(11); and the polymerase inhibitor Favipiravir (Figure 3D). We also performed a neutralization assay with an antibody raised against HNoV VLPs (Figure 3D). We could detect a significant reduction in HNoV replication in the case of nitazoxanide, TZX and Rupintrivir. Higher concentrations of Favipiravir could potentially show an antiviral effect, as no toxicity was observed at the concentration tested. Altogether, these data clearly demonstrate that 3D-HIE are amenable to infection with HNoV and to medium-throughput screening of anti-infectives.

### Host response signature upon infection suggest a functional activation of enteroendocrine cells

To evaluate the relevance of this system in addressing fundamental question about the host response against HNoV infection, we performed a population-wide RNA sequencing experiment on HNoV-infected and uninfected 3D-HIE. The experiment was repeated independently three times and the efficacy of infection was evaluated by RT-qPCR before sample submission (data not shown). Because of the preponderant interferon (IFN) signature that was detected in previous RNAseq dataset(9, 12), we included ruxolitinib to enrich for less represented, virus-associated pathways. First, we confirmed infection by detecting virus sequences in the HNoV-infected samples. We found 72984, 293942 and 48704 viral reads in the infected samples compared to 0, 2, 6 reads in uninfected samples, respectively (Figure 4A). In addition, we performed a multidimensional scaling analysis on the biological replicates that highlighted great variability between replicates and co-clustering of infected and uninfected samples from each independent experiment (RC4-RC1, RC5-RC2 and RC6-RC3, Figure 4B). For this reason, we pairwise analyzed the differentially expressed genes (DEG) in infected *versus* uninfected 3D-HIE (Volcano plot of DEG in Figure 4C). Due to the ruxolitinib treatment, IFN-stimulated genes were not differentially expressed between HNoV-infected and uninfected samples, although some IFN genes were upregulated upon infection (IFNA14 with fold change (FC)=5, IFNW1 with FC=4 and IFNL3 and L4 with FC of 2.3 and 1.8, respectively). A pathway analysis was next performed on significatively (p-value <0.05) down- and up-regulated genes (FC <-1.5 and >1.5, respectively) after removing genes with unknown codifying product (Table 1). Only 65 genes were significantly downregulated upon infection, and they strongly clustered in anti-inflammatory pathways, suggesting that the virus-induced innate immune response is mostly driven by bystander cells. On the other hand, over 500 genes were significantly upregulated in response to infection, but the pathway analysis did not highlight statistically significant signatures. However, the top upregulated pathways were olfactory transduction and response to hormones and neurotransmitters (dopamine, relaxine). Intriguingly, hormone and neurotransmitter receptors are mostly expressed on enteroendocrine cells(13) that have been recently described as a target of HNoV infection(14).

**Figure 4:**
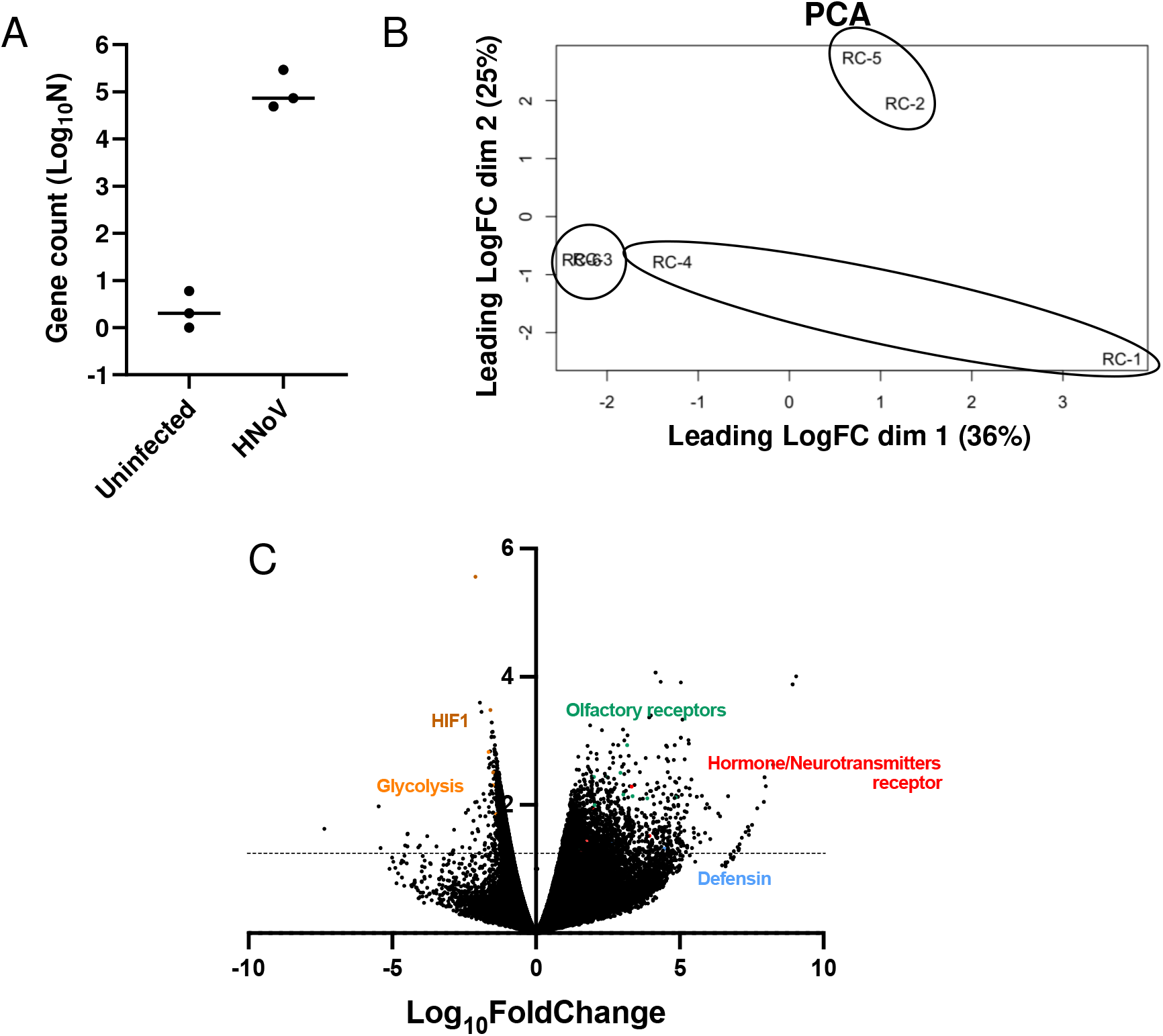
RNA sequencing of HNoV-infected 3D-HIE. 3D-HIE were infected with filter stool positive for GII.4 (#20942) for 2hrs at 37°C or kept uninfected. 3D-HIE were then washed twice in PBS, seeded in BME and cultured for 2d in the presence of GCDCA (500µM) and ruxolitinib (2µM). RNA was extracted with Direct-Zol RNA mini-kit and after quality control, the RNA was sequenced with the Illumina Hi-Seq 2500 platform. A) Viral reads were aligned to a consensus GII.4 Sydney (GenBank accession no. JX459908.1) and the counts were plotted in Graph Pad Prism. Each dot represents a biological replicate. B) Principal component analysis was performed to compare the biological replicates. Circles represents the independent repeats with the infected samples RC4, RC5 and RC-6 with the corresponding uninfected RC1, RC2 and RC3. C) Volcano plot of differentially expressed genes. On the y-axis, it’s the -Log_10_ p-value. Above the dotted line lay genes with significant p-value (p-value < 0.05).

**Table 1:** RNA sequencing pathway analysis.

|  | Name | P-value | Adjusted p-value | Genes |
| --- | --- | --- | --- | --- |
| Upregulated<br>(575 genes) | Olfactory transduction | 0.0003459 | 0.09513 | OR2C1, OR4Q2, OR8U1, OR2S2, OR2AE1, OR2W3, OR10W1, OR2AG2, OR11H6, OR56B1, OR8B4, OR5AP2, OR5B2, OR52B6, OR4D2, OR2B2, OR2T27, OR6J1, OR6N1, OR5P3, OR5P2, OR10A7, OR4K15, OR9A4, OR51T1, OR52A5 |
|  | Hormone ligand-binding G-protein coupled receptors | 0.003283 | 0.4514 | CGA, FSHR, GNRH2 |
|  | Signaling by GPCR | 0.006266 | 0.5355 | CXCL6, OR2C1, DHH, OR4Q2, RXFP2, PIK3R6, OR8U1, OR2S2, OR2AE1, OR2W3, OR10W1, NPY, OR2AG2, OR56B1, OR8B4, OR5AP2, CGA, HCRT, DRD3, OR5B2, WNT2, INSL5, OR52B6, DRD5, NTSR2, CCR2, OR4D2, OR2B2, OR2T27, WNT10A, OR6J1, TAAR5, OR6N1, OR5P3, OR5P2, OR10A7, OR4K15, OR9A4, FSHR, OR51T1, GNRH2, OR52A5 |
|  | Dopamine receptors | 0.007789 | 0.5355 | DRD5, DRD3 |
|  | Relaxin receptors | 0.02059 | 1 | RXFP2, INSL5 |
|  | Defensins | 0.03982 | 1 | DEFB121, DEFB124, DEFA6 |
| Downregulated<br>(65 genes) | HIF-1 transcriptional activity in hypoxia | 2.377E-06 | 0.0004137 | BHLHE40, SLC2A1, ADM, CA9, HK2 |
|  | BDNF signaling pathway | 2.171E-05 | 0.001889 | FOXC1, P4HA1, GDF15, IGFBP3, ADM, HSPA1B, HSPA1A |
|  | Apoptosis modulation by HSP70 | 0.001716 | 0.05855 | HSPA1B, HSPA1A |
|  | Glycolysis and gluconeogenesis | 0.001872 | 0.05855 | SLC2A1, PCK1, HK2 |
|  | Transcriptional regulation of white adipocyte differentiation | 0.002019 | 0.05855 | CEBPB, ANGPTL4, PCK1 |
|  | p53 signaling pathway | 0.001089 | 0.05855 | BNIP3L, GDF15, IGFBP3, HSPA1B |

Altogether, our data show that HNoV infection of 3D-HIE is a simplified and robust alternative to the established 2D protocol that is suitable for screening of anti-infectives and amenable to tuned virus-host interaction studies.

## Discussion

After 50 years from its discovery, there are still many knowledge gaps on HNoV tropism, biology and pathophysiology. Similar to other strictly enteric viruses, the lack of *in vitro* cultivation systems has also hampered the development of successful therapeutics to combat these prevalent virus infections. The discovery of HNoV replication in human intestinal enteroids in 2016(3) opened new avenues for norovirus research. However, limitations of this model exist in terms of labor, costs, and the lack of generation of a HNoV cell-derived stock. We sought to explore the possibility of simplifying the current protocol of infection of HIE to overcome some of the limitations. In our hands, the dispersion of 3D-HIE in 2D-monolayer prior differentiation has always been a variable step for a successful infection. The data presented herein demonstrates that HIE dispersion is not necessary for a successful HIE differentiation and that 3D-HIE spontaneously revert polarity upon differentiation, even if they are kept in BME. This finding was puzzling as a recent report suggests that HIE polarity is dependent on % ECM proteins(5). We reasoned that in our protocol, the 3D-HIE are kept in BME for more than the canonical 6 days and with every medium change, the concentration of ECM proteins could hence decrease. However, we also observed that in our model, terminal differentiation exacerbates the Apical-Out phenotype. Whether the observed spontaneous polarity is age and/or donor dependent deserves further studies. However successful infection (that can be used as proxy for successful differentiation and polarity inversion) was also observed in the adult jejunum line J2, albeit at lower efficacy, suggesting other factors also play a role in driving polarity inversion.

We successfully established infection of differentiated 3D-HIE with different HNoV GII.4-positive stool samples and across three institutions (Wobus lab at the University of Michigan, Rocha-Pereira lab at the KU Leuven and Taube lab at the University of Lübeck). As previously reported(9), ruxolitinib, a JAK inhibitor increased HNoV infection while the nucleoside inhibitors 2CMC reduced infection, corroborating that HNoV actively replicates in this model. Interestingly, the inversion of polarity alone as with the Apical-Out HIE did not ensure a successful infection by HNoV, suggesting that the BME provides an environment that is beneficial for a sustained viral infection.

We further demonstrated that this model is amenable to antiviral studies as we successfully reported activity of compounds that are endowed with antiviral activity against HNoV, such as Nitazoxanide and its prodrug TZX(10), and the protease inhibitor Rupintrivir(11). The model has also been successfully implemented for neuralization studies with an antibody raised against HNoV GII.4 VLPs, proprietary of R-biopharm.

Last, we wanted to explore whether this model would also provide insights in viral-host interactions and to this end, we performed an RNA sequencing analysis. Previous studies on enteric virus infection of enteroids unequivocally showed a signature dominated by interferon and innate immune pathways(7). Usually, interferon responses are initiated by infected cells but mostly amplified by bystander cells(6). In order to focus our analysis on signatures and pathways in infected cells as a consequence of viral replication, we included ruxolitinib, an inhibitor of JAK 1/2 and the downstream STAT-1 signaling, that limits interferon signaling. As a consequence, genes under the control of STAT-1 that might be upregulated upon infection will be reduced in expression. To our surprise, amongst downregulated pathways, we found HIF-1*α* pathways and glycolysis, that we previously described as an important host factor in the context of infection with the surrogate murine norovirus in macrophages(15). It is hence intriguing to wonder whether the previously observed “viral-induced” responses derive from actively replicating cells or they are a consequence of innate immune activation of bystander uninfected cells. Other possibilities are that macrophages and epithelial cells initiate different metabolically responses to infection, since mature enterocytes exhibit low rates of glycolysis(16) or that inhibition of the inflammatory response by ruxolitinib suppresses the inflammation driven glycolytic reprogramming(17). Studies that dissect the role and the contribution of uninfected bystander cells in an infected sample by single cell RNA sequencing are hence urgently needed.

Our RNA sequencing analysis could also shed some light into norovirus mechanisms of pathogenesis. Unlike the recently reported upregulation aquaporin (AQ) 1 upon HNoV infection of enteroids and potential involvement in HNoV-associated pathogenesis(18), we did not detect upregulation of AQ1 or other member of the AQ family in the RNAseq analysis. Instead, amongst the top upregulated pathways in response to infection, we found olfactory receptors and neurotransmitters signal transduction. Recently, human norovirus antigen was detected in enteroendocrine (EEC, Chromogranin A positive) cells from infected intestinal biopsies(14). EEC are rare cells in the gut epithelia (∼1%) and are chemosensors(13), hormone-producing cells in response of nutrients metabolites, bacteria metabolites, gut hormones and neurotransmitters. A subtype of EEC, the enterochromaffin cells have been recently described to be electrically excitable, and able to modulate the serotonin-sensitive primary afferent nerve fibers via synaptic connections and directly affect gut motility(19). Our analysis suggests a mechanism of viral-induced diarrhea by direct infection or by functional activation of enterochromaffin cells. It is also interesting to notice that the pathways positively associated to infection were not strongly upregulated suggesting that the signal from the rare EEC might be diluted in a bulk RNAseq analysis.

In summary, we propose here an adapted protocol for HIE infection by HNoV. This model is less time consuming, less costly, more reproducible, and more scalable for medium-throughput studies of antiviral/vaccine efficacy. In addition, because of the spontaneous polarity inversion, this model is amenable to infection by enteric viruses and likely other microbes without the burden of microinjection. We demonstrated the utility of this model for antiviral testing and studies of virus-host interactions. This model could also be useful for complex studies of HNoV-microbiome interactions.

## Material and methods

### Cells, treatment, and virus

Human intestinal enteroids derived from fetal ileum, HT124 and from adult jejunum, J2 were maintained in BME (MatriGel, Corning, 354234 or Cultrex Ultimatrix, BME001-05) and maintance media, CMGF+ (Advanced DMEM-F12, LWRN-conditioned medium, B27, N2, mrEGF, N-acetyl cysteine, Leu15-Gastrin, A-83-01 and SB202190), that is replaced every other day. Differentiation was triggered by Wnt3a removal with differentiation media (Advanced DMEM-F12, Noggin, B27, N2, mrEGF, N-acetyl cysteine, Leu15-Gastrin and A-83-01) for 6 days with medium change every other day. In the case of 3D-HIE, cells were kept in BME and treated with differentiation media 3-4 days post splitting, whereas in the case of 2D-HIE, cells were first subjected to trypsin treatment to create a cell suspension and then, they were seeded in collagen-coated 96-wells (10^5^ cells/well). The apical-out HIE were prepared according to the published protocol(20). Stool samples positive for HNoV GII.4 were prepared as a 10% solution in OptiMEM or PBS +/+ and then filtered through a 0.22µm membrane. The filtrate was then titrated by RT-qPCR, according to the published protocol(21), and used for infection at selected multiplicity of infection (MOI).

### Quantification of markers of differentiation by RT-qPCR

After 6 days of differentiation by Wnt3a withdrawal from the culture media, 2D- and 3D-HIE were harvested in TRI Reagent (Zymo Research, R2050-1) and cellular mRNA was extracted with DirectZol RNA extraction kit (Zymo Reseaerch, R2051). Two step RT-PCR was used to quantify relative expressions of intestinal cells markers (Sucrase Isomaltase for mature enterocytes, Mucin 2 for goblet cells, Lgr5 for stem-cells and Lysozyme for Paneth cells). The iScript cDNA synthesis kit (Bio-Rad) was used for cDNA synthesis. Quantitative real-time PCR (qPCR) with specific primers (Sucrase Isomaltase [SI]: forward AATCCTTTTGGCATCCAGATT and reverse GCAGCCAAGAATCCCAAAT; Lgr5: forward CAGCGTCTTCACCTCCTACC and reverse TGGGAATGTATGTCAGAGCG; Mucin 2 [MUC2]: forward TGTAGGCATCGCTCTTCTCA and reverse GACACCATCTACCTCACCCG; Lysozyme: forward ACAAGCTACAGCATCAGCGA and reverse GTAATGATGGCAAAACCCCA; GAPDH: forward CTCTGCTCCTCCTGTTCGAC and reverse TTAAAAGCAGCCCTGGTGAC) was performed with thermal cycler (Bio-Rad) by using iTaq Universal SYBR green supermix (Bio-Rad) as previously described(8). GAPDH was used to normalize gene expression.

### Immune fluorescence and image analysis

3D-HIE in maintenance media and differentiation media (both 10 days after splitting) were collected and washed from BME. In suspension, 3D-HIE were fixed with 4% paraformaldehyde for 30 minutes and then stained with DAPI to identify nuclei, and *Ulex Europeus* Agglutinin-1 (UEA-1) to label the villus brush border at the apical surface of the enterocytes. After staining, for ease of experimentation, 10-20 3D-spheres were transferred in a poly L-lysine coated black plate and subjected to automated high-content confocal imaging (CellVoyager, Yokogawa). Briefly, 16-maximum projection fields per well, covering more than 90% of the surface of the well were acquired with 10X objective. A Cell Profiler pipeline for image segmentation and a classifier for 3D-HIE UEA-1 positive and negative was then developed. The percentage of UEA-1 positive was obtained for each field and averaged for each well (technical replicate).

### Infection and viral quantification by RT-qPCR

HIE derived from fetal ileum, HT124, were maintained in BME (MatriGel, Corning, 354234 or Cultrex Ultimatrix, BME001-05) and in maintenance media for 3-4 days, then differentiated by Wnt3a withdrawal (differentiation media) for 5-6 days. On the day of infection, HIE were collected in a 15mL falcon tube and BME was washed with CMGF-medium. After pelleting (100g for 3 minutes at 4°C), HIE were transferred in a 1.5mL tube and the HNoV clinical isolate was added to selected MOI, according to the experiment. We estimated the MOI based on the number of spheroids per condition with an average of 300 cells/spheroid, as measured by immune fluorescence. Virus and cells were incubated in infection media (differentiation media and GCDCA, 500µM) for 2h at 37°C with occasional mixing. Then, HIE were washed 3 times with CMGF-, the 2h condition was harvested in TRI Reagent (Zymo Research, R2050-1) and the rest of infected-HIE were resuspended in BME and plated in a pre-warmed 24-well plate (3x 10uL drop/condition). Plates were left for 5 minutes at 37°C to ensure BME polymerization and infection media with ruxolitinib (2µM) and selected drugs were added. At 2dpi, cells were harvested in TRI Reagent and RNA extraction (Zymo Research, R2051) was performed according to manufacturer’s instruction. Viral quantification was performed as previously described(21).

### RNA sequencing and statistical analysis

HT124 were infected at MOI 1 or kept uninfected for 2d in the presence of GCDCA (500µM) and ruxolitinib (2µM). The infection experiment was performed three times. RNA was isolated using the DirectZol RNA extraction kit (Zymo Reseaerch, R2051). RNA library preparation and RNA-sequencing (single-end, 50 bp read length) were performed by the University of Michigan DNA Sequencing Core using the Illumina Hi-Seq 2500 platform. All RNA sequences were deposited in the NCBI GEO database and are cataloged under the accession number GSE205007. RNA sequences were generated for 151bp paired-end reads according to the manufacturer’s protocol (Illumina NovaSeq) by the University of Michigan Advanced Genomics Core. Bcl2fastq2 Conversion Software was used to generate de-multiplexed Fastq files. Raw sequences were mapped to a combined human genome GRCh38 (ENSEMBL) and the human norovirus GII.4 Sydney 2012 strain genome (GenBank accession no. JX459908.1), using STAR v2.5.2a(22). Aligned reads were then assigned count estimates to genes using featureCounts(23). Sample normalization and differential gene expression of the aligned sequences and all other analyses were calculated using the R package, EdgeR(24). We performed unbiased clustering of the normalized read counts using multidimensional scaling. To test for differentially expressed genes between viral-infected HIEs and mock-infected HIEs, by biological replicate, we applied the quasi-likelihood F-test using the glmQLFtest() function, controlling for biological replicates as a covatiate in analysis. The KEGG pathway enrichment analysis was performed using the kegga() function. Plots were constructed using the package ggplot2. Gene pathways were analyzed with the software Enrichr(25).

## Acknowledgements

CM is supported by Marie-Skłodowska Curie actions global fellowship (GA-841247). NSF is supported by an Horizon 2020-funded ITN program, OrganoVIR (Grant 812673). MG and JAC were supported by the National Institutes of Health (P30ES017885 and P30DK034933). We thank R-biopharm for graciously sharing anti-GII.4 antibody. We thank Jana Van Dycke for her support and fruitful discussions.

